# Mice in the Robbers Cave: Induction of intergroup conflict in mice using the competitive Tsunahiki task

**DOI:** 10.64898/2026.08.09.743721

**Authors:** Mariko Nakata, Nanako Fukai, Rino Iwabuchi, Haruto Muroyama, Joseph Carson, Yuen Yi Pun

## Abstract

Intergroup conflict is one of the most significant issues in human society. In the 1950s, Sherif et al. reported that intergroup conflict could be artificially induced in boys through intergroup competition with tug-of-war and ball games. Since this iconic study, researchers have developed various experimental methods to replicate intergroup competition and/or conflicts. However, although intergroup conflicts in wild animals are often reported, it has been difficult to establish a situation of intergroup conflict in laboratory rodents that is discriminable from aggressive behavior individually. In this study, we established a novel experimental paradigm for intergroup competition in mice in which the members of each group shared objectives and tasks. Adult male ICR/Jcl mice were housed in groups of six, divided into two teams of three and repeatedly performed a competitive Tsunahiki task (tsunahiki means tug-of-war in Japanese). The competitive Tsunahiki task was conducted in an open field divided into two experimental fields, with three ropes stuck to a wall separating the fields. The mice were required to pull two ropes out faster than their opponent team to win, and only the winners could proceed to the reward area separated by a guillotine door. We demonstrated that the experience of the competitive Tsunahiki task induced attack bites selectively toward members of the other team (out-group members). Our findings suggest that intergroup competition induces intergroup conflict in mice, providing a technical breakthrough in elucidating the detailed neuroscientific mechanisms underlying intergroup conflict.

## Introduction

War is not a privilege reserved for human beings. Wild animals, including chimpanzees (1) and meerkats (2) similarly have intergroup competition for resources. However, in laboratory rodents, a paradigm for inducing intergroup competition has not yet been established.

Researchers have attempted to observe conflict and reconciliation between human groups. One landmark study in social psychology, called “the Robbers Cave Experiment,” in which Sherif et al. (3) divided boys into two groups and made these groups compete with games, including tug-of-war. The experience of intergroup competition induces adversary behaviors toward out-group members belonging to the other group. Subsequently, the two groups were required to cooperate, which reduced intergroup conflict.

In the present study, we attempted to reproduce the Robbers Cave Experiment using mice. We divided the mice into two subgroups and experimentally induced conflict and reconciliation between the subgroups.

## Results and Discussion

Adult male ICR/Jcl mice (n = 84) were housed in groups of six (14 groups). After ten days of cohabitation, the formation phase was started (Figure 1A, left panel). In this phase, each group performed a group-based operant task: the Tsunahiki task for a single group (the single-group task) (4) (Figure 1B, left panel). In this phase, mice performed the task in a group, with one trial per day over five days in this phase. In this task, the mice were required to pull ropes placed in holes on the wall of the start area, which was one of the two areas divided by the guillotine door of an open field box. After all the ropes were pulled out, the door was opened and the mice could freely explore the reward area. Any individual may pull the ropes, and all group members can enter the reward area, regardless of their contribution. These rules created a situation in which all group members shared the same objectives and tasks.

**Figure 1.**
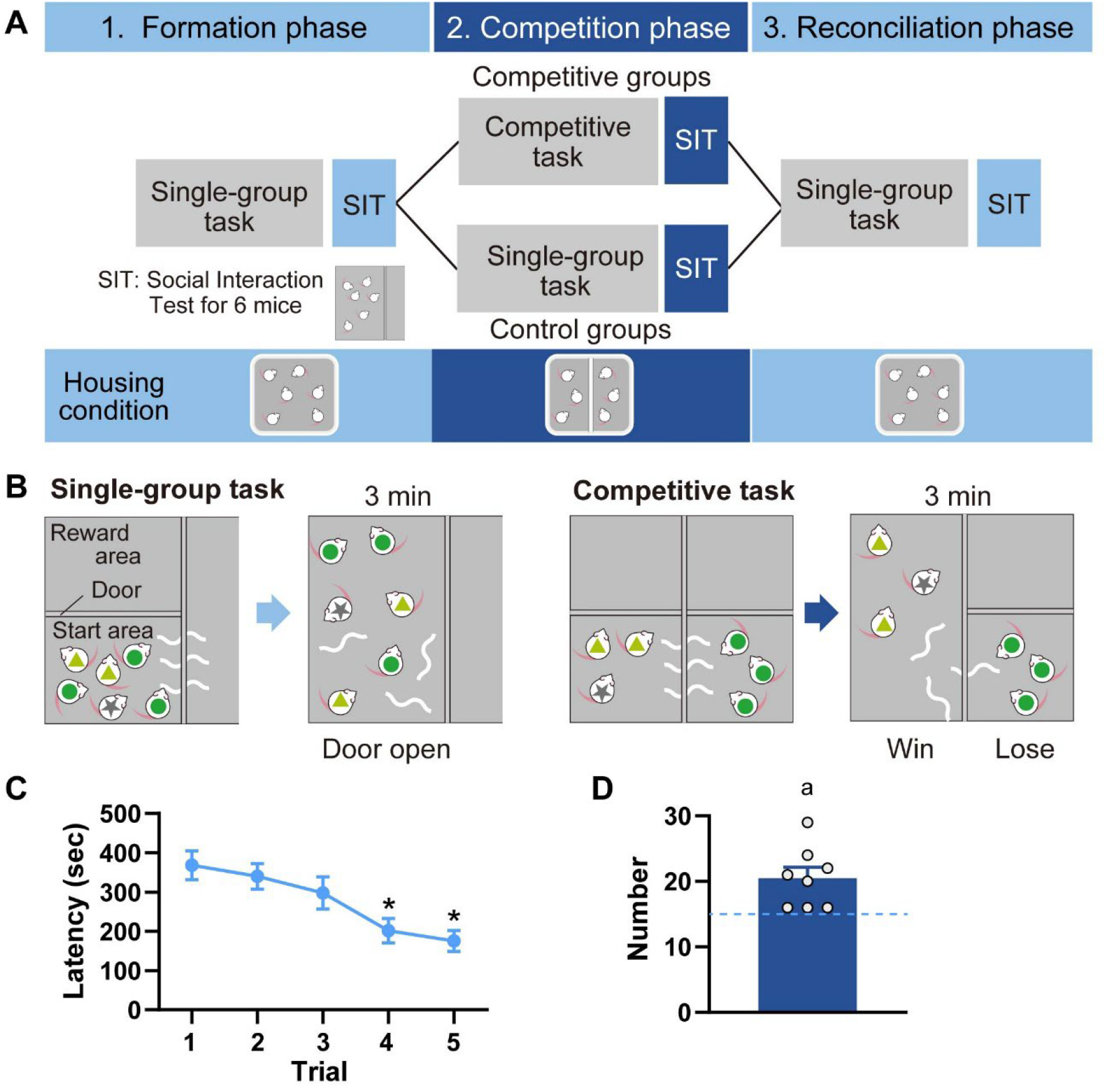
(A) Schematic diagram showing the flow of the entire experiment. Two teams were kept separately in the competition phase in both conditions and competed through the competitive task only in the competitive condition. (B) Schematic diagram showing the flow of the single-group (left panel) and the competitive (right panel) task in the Tsunahiki task. Mice markings with light green triangles and green circles indicate in-group (same team) and out-group (different team) members of a mouse marked with a gray star, respectively. (C) Latency to all ropes out in the single group trials during the formation phase (n = 14 groups). *: *p* < 0.05 vs trials 1 and 2. (D) Total number of wins in the 30 competitive trials during the Competition phase. The graph includes 8 subgroups, which won more frequently than their opponent subgroups during the competition phase. The dashed line indicates 15 trials, that is half of 30 trials. a: *p* < 0.05 vs 15. All data are shown as mean ± SEM, standard error of the mean.

All groups acquired the task in five trials with a significant reduction in latency to remove all ropes (Figure 1C, *F*_(4,52)_ = 11.211, *p* < 0.001, *η*^*2*^ = 0.275) and then proceeded to the competition phase (Figure 1A, middle panel). At the beginning of this phase, each group was divided into two teams of three mice each. The two teams were kept cohabiting in their home cages, but a transparent and perforated partition was introduced to separate the teams with contact. All groups underwent three trials per day for ten days during this phase. Eight groups were assigned to the competitive condition and repeatedly experienced the competitive Tsunahiki task (Figure 1B, right panel; Movie S1). The two teams faced each other across a transparent and perforated wall and competed to pull the ropes. A team pulling two ropes first won and explored the reward area freely for three minutes. However, the losing team experienced delays in the start area. No team won all 30 trials, but the number of wins by the teams that won more frequently than their counterparts significantly exceeded 15, half of all trials (Figure 1D, *t*_(7)_ = 3.383, *p* = 0.012, vs 15, *d* = 1.196). The remaining six groups were assigned to the control condition. They were housed in partitions similar to those of the competitive groups; however, the two teams continued the single-group task together. After the end of the competition phase, the groups underwent a reconciliation phase in which all groups were required to execute the single-group task for one trial per day for six or seven days (Figure 1A, right panel). Partitions in the home cages were removed at the beginning of this phase. At the end of each phase, all groups underwent the social interaction test (SIT), in which the mice were allowed to explore the experimental field freely with the two teams mixed, and aggressive behavior during the SIT was observed.

The control groups did not show any significant differences in the number of attack bites in the SIT among the three trials and between the in-group and out-group targets, that is, individuals in the same and different teams in the competitive phase (Figure 2A, left panel). Surprisingly, however, groups in the competitive condition responded remarkably to the competitive Tsunahiki task. All groups began to show attack bites selectively toward out-group members only in the second trial of the SIT, immediately after the competition phase. In contrast to Trial 2, almost no attack bites were observed in Trials 1 or 3 after the formation and reconciliation phases (Figure 2A, right panel; Target: *F*_(1, 7)_ = 9.952, *p* = 0.016, *η* ^*2*^ = 0.098, Phase: *F*_(1.03, 7.19)_ = 16.881, *p* = 0.004, *η* ^*2*^ = 0.319, Interaction: *F*_(1.02, 7.11)_ = 11.879, *p* = 0.010 *η* ^*2*^ = 0.210). In teams from the competitive groups, the total number of bites in Trial 2 was significantly and positively correlated with the number of wins in six competitive trials in the last two days before SIT (Figure 2B, *rho* = 0.663, *p* = 0.005). This result indicates that the winning team members tend to attack the losing team members more. Moreover, most of the 15 attackers in the competitive groups showed attack bites only in Trial 2 (Figure 2C, right panel), but this was not the case in the control group (Figure 2C, left panel, rate of “only phase 2” attackers: *p* = 0.041, OR: 7.282 between conditions). These results suggest that intergroup competition induces attack bites even in individuals with a low baseline aggression.

**Figure 2.**
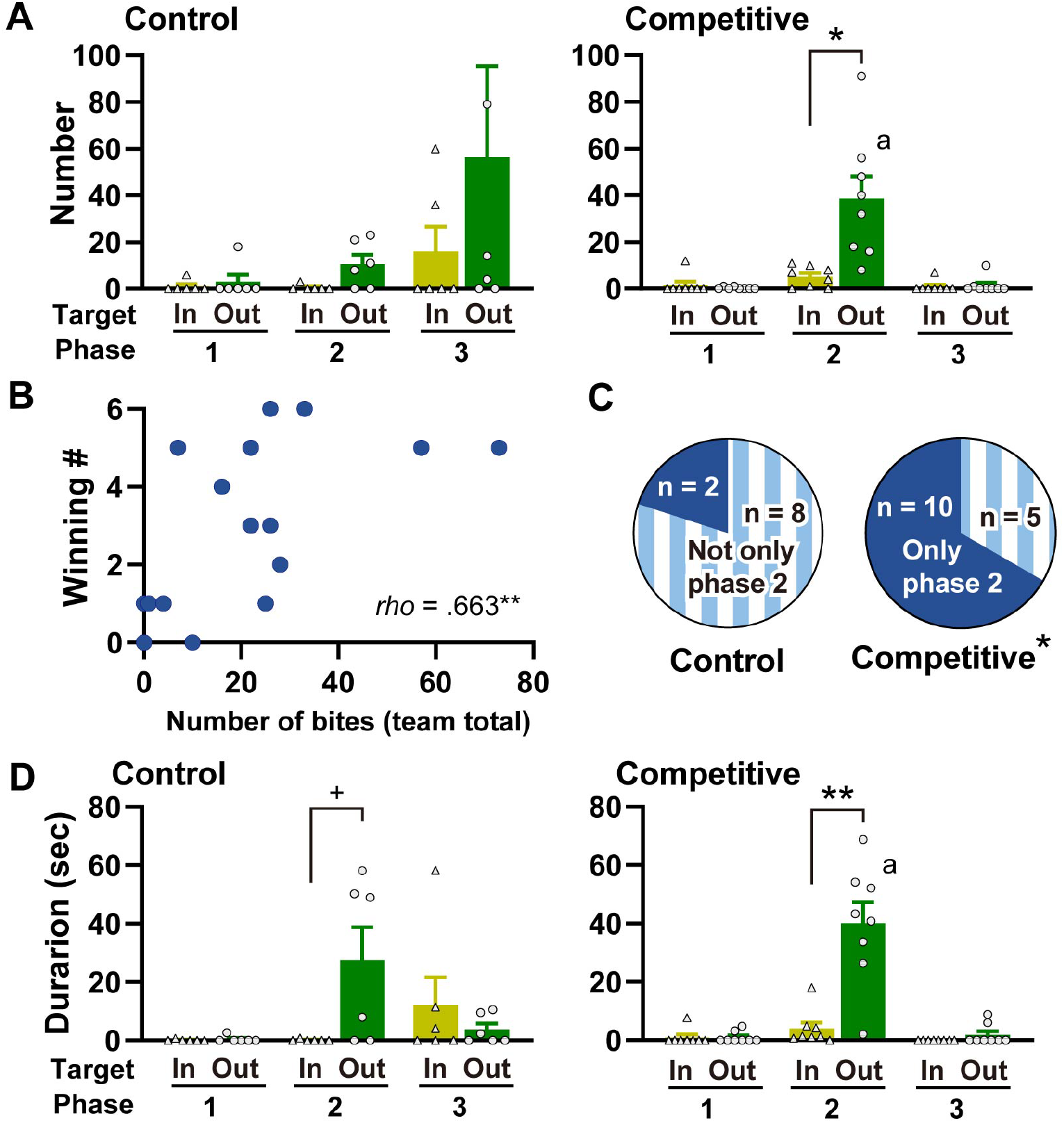
(A) Total number of attack bites during each trial of the SIT in the control (left panel) and competitive (right panel) conditions. Light green bars with the target label “In” indicate number of attack bites toward in-group members. Deep green bars with the target label “Out” indicate number of attack bites toward out-group members. *: *p* < 0.05; a: *p* < 0.05 vs phase 1 and 3 of the same targets. (B) Relationship between total number of wins in the last 6 trials before the SIT and total number of attack bites in the second SIT after the competitive phase in each team. *rho*: correlation coefficient, **: *p* < 0.01. (C) Number of mice that showed attack bites only in the competitive phase (phase 2) (solid deep blue) and those also showed attack bites in the formation and/or reconciliation phases (not only phase 2) (striped light blue). Animals that showed at least one attack bite throughout the three SIT trials were included. *: *p* < 0.05 test vs control. (D) Total duration of sideways threats and pursuits (SWP) during each trail of the SIT under the control (left panel) and competitive (right panel) conditions. Light green bars with target label of “In” indicate duration of SWP toward in-group members. Deep green bars with target label of “Out” indicate duration of SWP toward out-group members +: *p* < 0.10, **: *p* < 0.01; b: *p* < 0.01 vs phase 1 and 3 of the same targets.

It should be noted that not only were the competitive groups statistically significant (Figure 2D, right panel; Target: *F*_(1, 7)_ = 20.783, *p* = 0.003, *η* ^*2*^ = 0.146. Phase: *F*_(1.03, 7.18)_ = 33.259, *p* < 0.001, *η* ^*2*^ = 0.358, Interaction: *F*_(1.04, 7.26)_ = 19.730, *p* = 0.003, *η* ^*2*^ = 0.250), but the control group tended (Figure 2D, left panel; Target: *F*_(1, 5)_ = 4.080, *p* = 0.099, *η* ^*2*^ = 0.036, Phase: *F*_(2, 10)_ = 3.229, *p* = 0.083, *η* ^*2*^ = 0.109, Interaction: *F*_(1.18, 5.92)_ = 4.380, *p* = 0.079, *η* ^*2*^ = 0.207) to show an out-group-targeted increase in cumulative durations of sideways threats and pursuits, which are milder forms of aggression than attack bites. Thus, in addition to the experience of competition, the separation of two teams in their home cages may have increased mild aggression selectively toward out-group members, similar to a previous report in male mice (5). These results collectively suggest that intergroup competition using the Tsunahiki task successfully induced conflict between teams, as indicated by the increased attack bites in male mice. Intergroup conflict is the seed of serious social problems, such as war and discrimination. Replicating this phenomenon in laboratory rodents would enable the establishment of biological interventions to address these issues.

## Materials and Methods

The experimental design is described in the results section. Detailed information on the materials and methods used is available in the Supplemental methods.

### Animals

Eighty-four adult male ICR/Jcl mice were divided into 14 groups (6 mice each) and marked with dye on their backs upon arrival. The mice were housed under a 12-h light–dark cycle (dark phase started at noon) and provided with food and water *ad libitum*. All behavioral tests were conducted in the dark phase under dim light (15 lx). All experiments were approved by the Animal Care and Use Committee of the University of Tsukuba and conducted according to NIH guidelines. All efforts were made to minimize the number of animals used and their suffering.

### Experimental apparatus

All behavioral experiments were conducted in a gray opaque acrylic open field box (90 cm × 90 cm × 40 cm), divided into two experimental fields using a transparent and perforated acrylic wall. Each field was further divided into two areas (start and reward areas for the Tsunahiki task) using a transparent and perforated guillotine door. The experiments were recorded using digital video cameras placed above the apparatus.

### Tsunahiki task

In brief, mice were placed in the start area and required to pull on ropes placed in holes in the dividing wall between the two experimental fields. When the ropes were pulled out as required, the guillotine door was manually opened, and the mice could freely explore the reward area. The procedures for the single-group task have been described previously (4), and the latency to the ropes being pulled out was recorded for each trial. The competitive task was newly developed and described in the results section. The team that won each trial was recorded.

### Social interaction test (SIT)

Six mice in each group (including the two teams) were released into an experimental field without a guillotine door. They were allowed to freely explore and interact for 20 min. All video recordings were scored by an experimenter using the BORIS digital event recorder program (6). Aggressive behavior was scored based on the method of a previous study (7).

### Tube test

Tube tests were conducted using previously described procedures (8, 9). Briefly, each experimental animal was trained to walk through a transparent acrylic tube of 30 cm length and 3 cm inner diameter for two training days. All the pairs within a group were tested once in a round-robin manner on each test day. The individual who stayed longer in the tube was identified as the winner of the pair. The rank of each individual was determined based on the number of wins.

### Statistical Analysis

Statistical analyses were performed using the R software (version 4.1.0) (10). Data were analyzed using one-way repeated-measures ANOVA, two-way repeated-measures ANOVA, one-sample t-test (two-tailed), Spearman’s correlation test, or Fisher’s exact test.

## Acknowledgments

We thank Dr. S. Yamamoto, Dr. S. Ogawa, Dr. T. Setogawa, Dr. S. Yamane, Dr. Y. Watanabe and R. Tamura for valuable discussions, and R. Akai, M. Kobayashi, H. Nakagawa for support for technical assistance. We also thank A. Sagehashi for secretarial assistance. English-language proofreading was conducted by Editage. This work was supported by grant-in-aid for Scientific Research (KAKENHI) 19K14491, 22H01099 and 23K22370 to MN.

