## Supplemental methods for "Mice in the Robbers Cave: Induction of intergroup conflict in mice using the competitive Tsunahiki task"

**This PDF file includes:**

Supporting text
Figures S1 to S7
Legends for Movies S1
SI References

**Other supporting materials for this manuscript include the following:**

Movies S1

### Supplemental methods

#### Animals and housing conditions

Adult male ICR/Jcl mice (n = 84, eight weeks old upon arrival) were purchased from a commercial breeder (CLEA Japan, Inc., Japan). On the day of arrival, mice were assigned to a group of six, each comprising individuals that had been transported in the same cage and were close in body weight. Average body weight was 36.7±1.09 g (mean±SD) and the variation in weight within a group was set within 2.2 g. On the same day, mice were anesthetized with isoflurane and their back fur was stained with black hair dye for individual identification, as described in Figure S1. As the markings of all individuals remained sufficiently distinct to allow for identification until the end of the experiment, there was no need to dye their fur again during the experimental period.

After the staining, mice were kept in groups in white plastic cages (36 × 36 × 17 cm) with white paper bedding (ALPHA-dri, Shepherd Specialty Papers, USA). A transparent acrylic tube 10-cm in length was introduced into each cage for habituation to the tubes used in the tube test. The tube was removed from the home cage on the first day of habituation to the experimental apparatus. The room temperature was maintained at 23±2°C and food and water were available *ad libitum* throughout the experiment. All behavioral tests were conducted in the dark phase of a 12-h light-dark cycle (the dark phase started at noon). For the behavioral tests, mice were moved to an experimental room for more than two hours before the test started. All behavioral tests were conducted in the dark phase under dim light (15 lx). All experiments were approved by the Animal Care and Use Committee of the University of Tsukuba and were conducted following NIH guidelines. All efforts were made to minimize the number of animals used and their suffering.

#### Experimental design

Experiments were conducted in two batches. Batch 1 included six groups and batch 2 included eight groups. For both batches, behavioral tests were conducted according to almost the same schedule (Figure S2). The entire experiment comprised three phases, as shown in Figure 1A and S3: the formation phase (phase 1), competition phase (phase 2), and reconciliation phase (phase 3).

Behavioral experiments were initiated 10 days following arrival. On the first and second day, the mice were habituated to the experimental apparatus and trained for tube testing. Training for the tube test was conducted after all groups had finished their habituation to the apparatus. Tube tests were conducted on the third day.

The formation phase (phase 1), starting from the fourth day, was intended to establish social relationships within each group of six mice and promote the acquisition of the single-group Tsunahiki task. Each group was trained for the single-group Tsunahiki task with one rope and all groups successfully pulled the rope out in one trial. Thereafter, each group performed the single-group Tsunahiki task for five trials, with one trial per day. At the end of this phase, the social rank within each group was determined using the tube test and the social interaction test (SIT) was conducted (Figures S2 and S3).

The competition phase, performed following the formation phase, examined the effects of intergroup competition using the Tsunahiki task on social interaction in the SIT. At the start of this phase, groups were assigned to either the control (six groups, two from batch 1 and four from batch 2) or competitive conditions (eight groups, four from batches 1 and 2 each). Six individuals in each group were divided into two teams of three mice each. Team members were determined based on the results of the second tube test, with one team comprising individuals ranked 1st, 3rd and 5th, and the other comprising those ranked 2nd, 4th and 6$^{th}$ to ensure that the social rank within each team was as uniform as possible. However, when a mouse repeatedly displayed aggressive behavior targeting a specific individual during the formation phase, the aggressor and target were assigned to different teams regardless of their social rank. This exceptional measure was necessary for only one group in batch 1.

In the competition phase, the groups in the competitive condition repeatedly experienced competitive Tsunahiki task trials, with three trials per day for ten days. Two teams from the same group always competed in a competitive task. In contrast, the control groups underwent the same number of trials as the single-group task, with six mice in each group (see Figure S3). After the end of the trials on the first day of the competition phase, transparent and perforated partitions (35.5 cm × 16.5 cm, Figure S4A) were introduced into the home cage of the groups in both conditions, and two teams were cohabited while separated by the partition (Figure S4B). Each team was assigned a specific position on either side of the partition, which was maintained throughout the competition phase. The partition was removable and contained nine holes, each 1 cm in diameter, allowing individuals from different teams to investigate each other. Both teams had free access to food and water. At the end of this phase, the social rank of each team is determined using a tube test. The SIT with six mice in each group was also conducted after the tube test.

The reconciliation phase examined whether the two teams could unite after the competition by executing a single-group task together. All the groups performed the single-group Tsunahiki task, similar to the formation phase, for seven (batch 1) or six (batch 2) trials, with one trial per day (see Figure S3). After the completion of the first trial in the reconciliation phase, each group was returned to its home cage without any partitioning. The tube test was conducted twice during the reconciliation phase after the first and sixth trials. At the end of this phase, SIT was conducted as in the other phases.

It should be noted that four mice (one mouse in each group from four groups) died during phase 2 of the experimental period. They died accidentally, and not because of aggressive behavior from their cage mates. These mice included one mouse from the control group (died on day 30 of batch 2; experimental days are indicated in Figure S2) and three mice from the competitive groups (died on day 34 of batch 1, days 23 and 27 of batch 2). Even after one team consisted of two mice, the experiment continued using the same procedure as the other groups. Deaths on day 34 in batch 1 and day 23 in batch 2 did not affect the competition results because they occurred after the end of all competitive trials (day 34) or after the end of the first day of the competitive phase. However, death on day 27 induced a decrease in the team from 85.71 (14 trials) to 53.33 (15 trials).

**Experimental apparatus**

All behavioral experiments were conducted in a gray opaque acrylic open field box (90 cm × 90 cm × 40 cm, LE800SC, Panlab, S.L.U., Spain), divided into two rectangular experimental fields (90 cm × 45 cm × 40 cm) by a transparent and perforated acrylic wall. The wall had 12 holes, each 1 cm in diameter, located 3 cm above the floor. These holes allowed the experimenter to place the ropes through them and allowed the mice to investigate the filed on the other side. The use of the two fields was counterbalanced by experimental day and group/team. Each field was further divided into two areas (start and reward areas for the Tsunahiki task) using a transparent and perforated guillotine door. The guillotine door had three holes, each 1 cm in diameter, and 3 cm above the floor. It was attached to a pulley at the top of the chamber using a clear nylon string, and the experimenter manually pulled the string to open the door. For the Tube test, the doors were closed and one of the areas was used. For the SIT, the door was removed and each rectangular field was used (Figure S5). The experimental apparatus was lit with white light (15 lx). The apparatus was covered with black curtains and the mice could not see the experimenter during the behavioral tests (Figure S5). At the end of each trial, the apparatus was wiped using 70 % ethanol. Experiments were recorded using digital video cameras placed above the experimental field.

**Habituation to the apparatus**

Prior to the formation phase, each group was habituated to the experimental apparatus. Mice were allowed to explore one of the experimental fields in groups of six, with the door open. Six mice were released from a black plastic starter box and returned to their home cages after 20 min of exploration. On the second day, the opposite side of the experimental field was used, allowing all groups to habituate to both fields.

**Tsunahiki task**

**Apparatus.** The Tsunahiki task used plastic strings 15 cm in length (ropes). Although the ropes were originally twisted, we opened them when they were placed in the holes to prevent them from falling owing to accidental contact with the mouse (Figure S6). The ropes were placed in holes in the wall separating two experimental fields. When the groups were trained using a single rope, the rope was placed in the third hole from the door. Three ropes for the usual trials of the single-group and competitive tasks were placed in the first, third, and fifth holes from the door. No pellets or food rewards were placed in the reward area because it has been previously demonstrated that entry into the reward area itself seems to be rewarding enough for mice (1).

**Procedures for the single-group task.** One experimental field was used for the single-group task. At the beginning of the task, mice were released from a black opaque plastic start box (14 × 14 × 12 cm) placed at the edge of the start area. When the mice pulled out all the ropes that were placed in the holes on the wall within 20 min, the guillotine door was manually opened by an experimenter who monitored the behavior of the animals through a video camera, and the mice could explore the reward area for three minutes. All mice entered the reward area regardless of which individual executed the task (Figure 1B, left panel).

If the rope accidentally fell into the opposite experimental field before being pulled out, the experimenter quickly returned the rope to its original position. If the rope fell off accidentally from the hole into the experimental field without active rope-pulling behavior by the mouse with its forelimbs or mouth, the experimenter did not intervene. The experimenter manually recorded the latency to all ropes being pulled out for each trial. All test trials were recorded using a digital video camera placed above the experimental field for detailed analysis.

**Procedures for the competitive task.** Two experimental fields were used for the competitive task. Each team of the same group was placed in the experimental field. Each team was simultaneously released from the same start box as in the single-group task placed at the edge of each start area. Three ropes were placed in the holes in the wall, and each team pulled the ropes into their own field. There were six holes larger than the number of ropes, and the mice were able to investigate the opposite experimental field from the empty holes.

The team that pulled out two of the three ropes first won the trial; an experimenter manually opened the door of the winning team, and then the team could explore their reward area freely for three minutes. The losing team had to wait during the exploration, that is, they experienced a three-minutes delay (see Figure 1B, right panel, and Figure S7). In the competitive task, the experimenters did not intervene in the accidental rope-outs. An accidental rope-out into the opposite experimental field was counted as a rope-out by the opposing team, which was similar to their own goal. All test trials were recorded using a digital video camera placed above the experimental field for detailed analysis.

**Tube test**

Tube tests were conducted as described previously (2, 3) to assess the rank of each mouse within a group or team. For rank determination within a group of six mice, the test was conducted 15 times per group in a round-robin tournament. For rank determination within a team of three mice, the test was conducted thrice per team in a round-robin tournament. A transparent acrylic tube (inner diameter: 3 cm, length: 30 cm) was placed in the experimental field for training and testing.

**Training.** Each mouse was trained to run through a tube for two consecutive days before the first tube test. The mice were individually placed in the tube and allowed to run back and forth four times per day. If the mouse attempted to exit the tube while backing up, the experimenter gently pushed it back into the tube. The training for the two round trips was conducted immediately before the test on each testing day.

**Test.** Two mice were placed at each end of the tube and subsequently released to start when the whole body of both mice entered the tube. When all limbs of one mouse emerged from the tube, the match ended. The individual that emerged from the tube was the loser (subordinate), and the individual that remained in the tube was the winner (dominant). If no mouse was ejected within two minutes, the trial was considered a tie, although no trial resulted in a tie. When a mouse refused to enter the tube, it was recorded as a loser (forfeited match).

**Behavioral analysis and rank determination.** The experimenters manually recorded the number of wins for each mouse, and the latency to the outcome was determined for each match. Generally, individuals with more wins within each group or team are ranked higher. If there were multiple individuals with the same number of wins, the result of the match(es) where the individuals in question faced each other directly was referred to, and the mouse that won directly ranked higher. In the present study, the ranks from the second tube test were used to determine the team members in the competition phase. At that time, one pair showed the same number of wins, but a direct match was not held because of refusal by both individuals. Because these two mice did not win at all compared to the other individuals, the mice with a longer cumulative latency to lose ranked higher than the other mice.

#### Social Interaction Test (SIT)

**Procedures.** The SIT was conducted thrice at the end of each phase using one experimental field without a door. In all three trials, six mice in each group were released together from the same start box as in the Tsunahiki task at the beginning and were allowed to explore and interact freely for 20 min. All test trials were recorded using a digital video camera placed above the experimental field to analyze aggressive behavior.

**Behavioral analysis.** The aggressive behavior of each individual during the SIT was analyzed using BORIS version 9.6.5. (4). Four types of aggressive behaviors (attack bites, sideways threats, pursuits, and tail rattles) were recorded based on a previous study (5). However, few tail rattles were observed, and tail rattles were not included in the data analysis. The targets of each aggressive behavior were also recorded. If aggressive behavior targeted a member of the same team, that behavior was defined as being directed toward an in-group member. Alternatively, an aggressive behavior targeting the members of a different team was defined as toward an out-group member. The total number of attack bites and the cumulative duration of sideways threats and pursuits of in-group or out-group members were calculated for each group or team.

#### Statistical Analysis

Statistical analyses were conducted using R software (version 4.1.0) (6). Latency to all ropes being pulled out was analyzed using one-way repeated-measures ANOVA, with trials as the within-subject factor. A post-hoc multiple comparison was conducted using the Holm correction. The number of wins was compared to half of all the trial numbers (15) using a one-sample t-test (two-tailed). Aggressive behaviors, including the total number of attack bites and cumulative durations of sideways threats and pursuits, were analyzed using a two-way repeated-measures ANOVA with phases and targets as the within-subject factors. When the assumption of sphericity was violated, the Greenhouse–Geisser correction was applied. Post hoc multiple comparisons were conducted using the Holm correction. The association between the number of wins and attack bites was analyzed using Spearman's rank correlation coefficient (*rho*). Differences in the proportion of individuals showing attack bites in the SIT only after phase 2 between the control and competitive conditions were assessed using Fisher's exact test. Differences were considered statistically significant at $p < 0.05$.

**Figures**

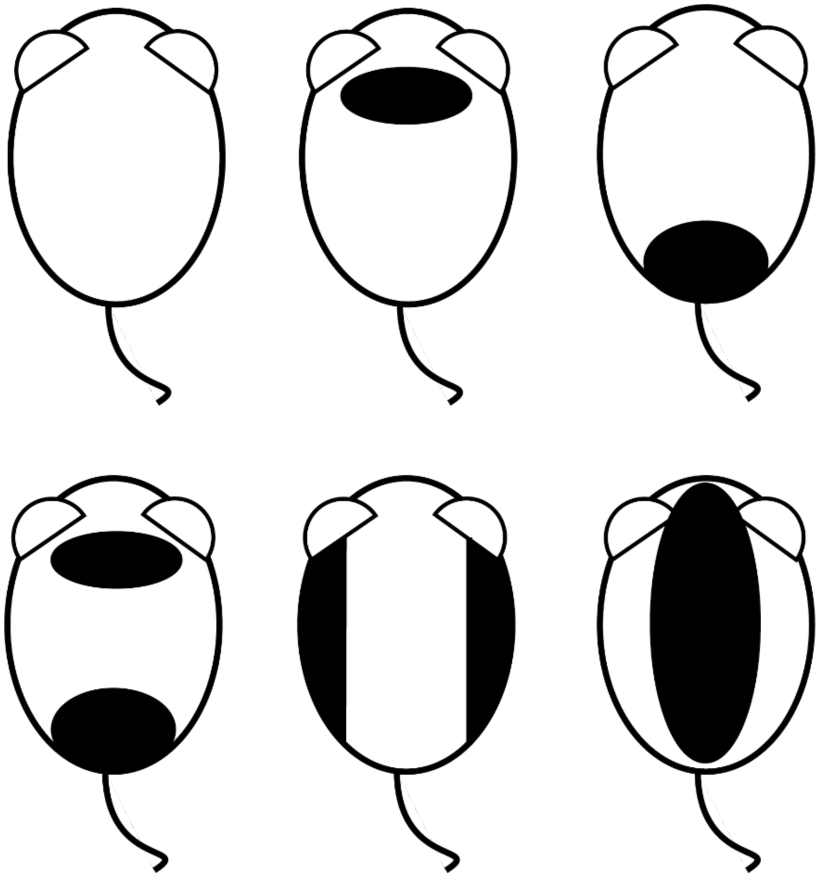

**Figure S1.** Schematic diagram of the fur dyeing pattern for individual identification in each group. The black part indicates the section stained with black hair dye. One mouse per group was left without staining (top left).

| | Batch1 | | Batch2 | |
|---|---|---|---|---|
| Day | Control | Competitive | Control | Competitive |
| 0 | Arrival | | Arrival | |
| 1 | | | | |
| 2 | | | | |
| 3 | | | | |
| 4 | | | | |
| 5 | | | | |
| 6 | | | | |
| 7 | | | | |
| 8 | | | | |
| 9 | | | | |
| 10 | Habituation + Tube training (2 days) | | Habituation + Tube training (2 days) | |
| 11 | | | | |
| 12 | | | Tube test (1) | |
| 13 | Tube test (1) | | Tsunahiki Training | |
| 14 | Tsunahiki Training | | Single-group task | |
| 15 | Single-group task | | | |
| 16 | | | | |
| 17 | | | | |
| 18 | | | | |
| 19 | | | | |
| 20 | Single-group task | | Tube test (2) | |
| 21 | Tube test (2) | | SIT (Phase 1) | |
| 22 | SIT (Phase 1) | | | |
| 23 | Single-group task | Competitive task | Single-group task | Competitive task |
| 24 | | | | |
| 25 | | | | |
| 26 | Single-group task | Competitive task | | |
| 27 | | | | |
| 28 | | | | |
| 29 | | | | |
| 30 | | | | |
| 31 | | | | |
| 32 | | | | |
| 33 | | | Tube test (3) | |
| 34 | Tube test (3) | | SIT (Phase 2) | |
| 35 | SIT (Phase 2) | | Single-group task | |
| 36 | Single-group task | | Tube test (4) | |
| 37 | Tube test (4) | | Single-group task | |
| 38 | Single-group task | | | |
| 39 | | | | |
| 40 | | | | |
| 41 | Single-group task | | Single-group task | |
| 42 | | | | |
| 43 | | | Tube test (5) | |
| 44 | Single-group task | | SIT (Phase 3) | |
| 45 | | | | |
| 46 | Tube test (5) | | | |
| 47 | SIT (Phase 3) | | | |

**Figure S2.** Detailed experimental schedule for batch 1 and batch 2. SIT: Social Interaction Test.

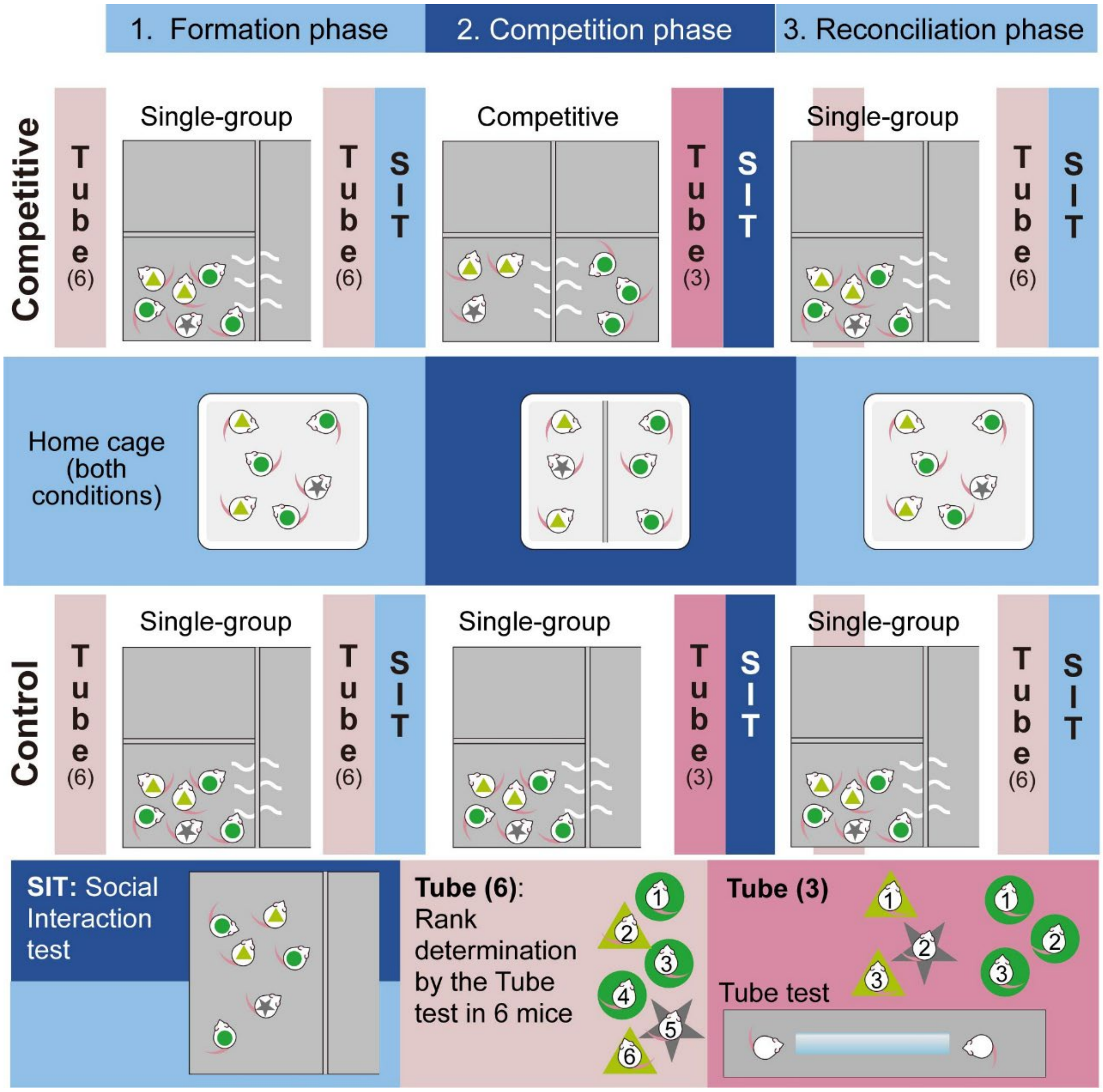


**Figure S3.** Schematic diagram of the detailed flow of the whole experiment. Mice marked with light green triangles and green circles indicate in-group (same subgroup) and out-group (different subgroup) members of a mouse marked with a gray star. The first row indicates the flow of the whole experiment in the competitive groups. The third row indicates the flow of whole experiment in the control groups. Pink bars indicate tube tests and blue bars indicate SITs. The second row indicates housing conditions in both competitive and control groups in three phases. The numbers on the mice in the bottom row indicate examples of social rank within a group or a team. SIT: Social interaction test, Tube: Tube test.

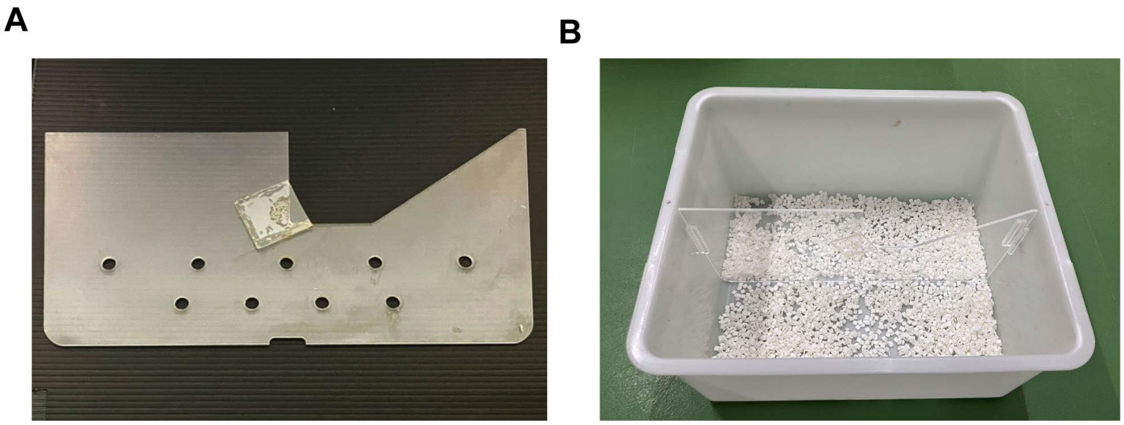

**Figure S4.** Photo of the home cage used in the competition phase. (A) Photo of the partition that separated two teams. (B) Photo of the home cage with the partition. The same cages were also used in the formation and reconciliation phases without the partition.

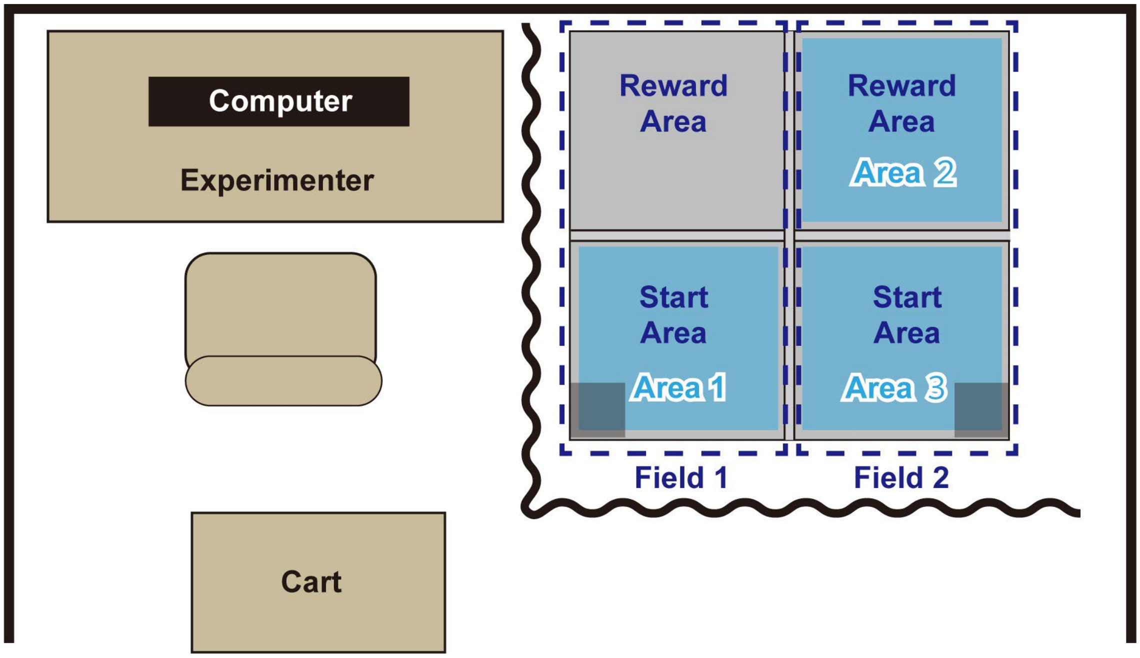


**Figure S5.** Experimental apparatus for the behavioral tests. Schematic diagram of the entire experimental booth used for the behavioral tests. Fields 1 and 2, surrounded by deep blue dashed lines, were used for the Tsunahiki task and the SIT. Start and reward areas for the Tsunahiki tasks were indicated by deep blue characters. The semi-transparent black squares at either end of the start area indicate the place of the start boxes. Three areas (Areas 1-3) were used for the Tube test with the guillotine doors closed. Black curtains were closed during the Tsunahiki task and the SIT, and mice could not see the experimenter sitting in front of the computer on the left side of the apparatus. Cages were placed outside of the experimental booth covered with black curtains and transferred onto the cart right before their first trial.

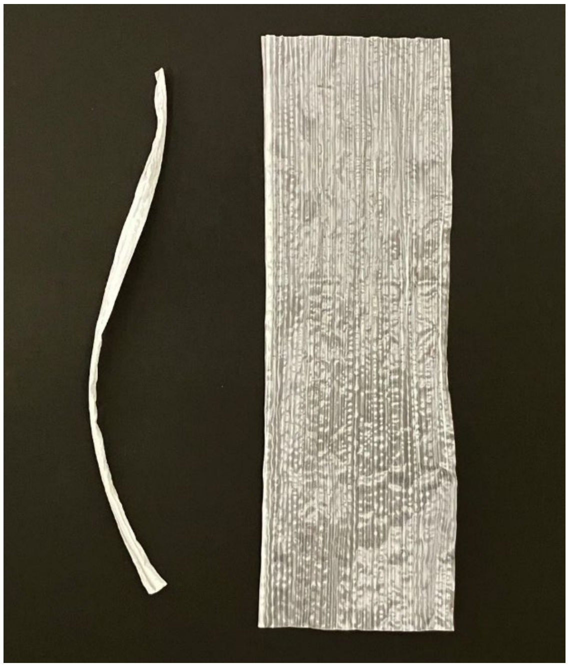

**Figure S6.** Photo of the ropes used in the Tsunahiki task. Left: twisted, right: opened.

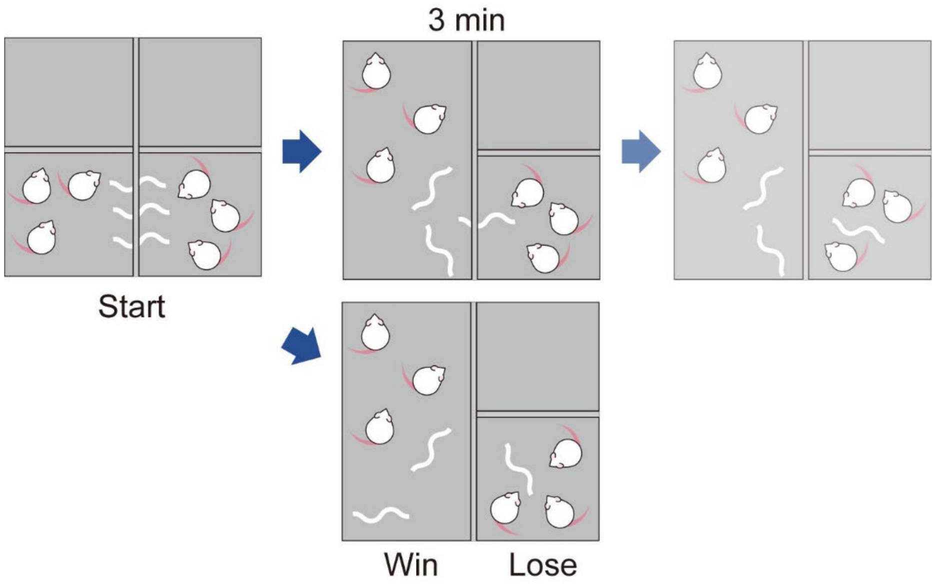


**Figure S7.** Schematic diagram of the detailed flow of the competitive Tsunahiki task. There were two types of matches. Sometimes one team pulled out two ropes initially (top row). Even in that case, pulling out of the remaining rope, often by the lost team, was observed after the door of the winning team opened. Alternatively, two teams pulled out one rope each, and then the last rope was pulled out by one, the winning team (bottom row).

**Movie S1 (separate file).** Representative video showing the performance in the competitive Tsunahiki task. This video is from batch 1 and recorded during the second trial of day 9.
